# Evaluation of acaricidal activity of lithium-chloride against *Tetranychus urticae (Acari: Tetranychidae)*

**DOI:** 10.1101/2022.07.19.500612

**Authors:** Izabella Solti, Éva Kolics, Sándor Keszthelyi, Zsuzsanna Bacsi, Ádám Staszny, Erzsébet Nagy, János Taller, Kinga Mátyás, Balázs Kolics

## Abstract

*Tetranychus urticae* is a severe threat and a major source of yield loss in some agricultural and horticultural crops. Control is greatly complicated due to the hidden lifestyle of the species. In addition to its direct damage, it is considered a vector for several viruses. The number of active substances used against it is limited, and the development of resistance means that sustainable control is not achieved. Therefore, there is a continuous need for new active substances. Lithium salts have been shown to be a promising alternative to control *Varroa destructor*, an apicultural mite pest. Therefore, in the present study, we investigated whether the contact efficacy of lithium chloride extends to other agricultural mite pests, such as the two-spotted spider mite. In the present pilot study, we report for the first time that the efficacy of lithium chloride extends to the two-spotted spider mite, showing 100% mortality at concentrations of 5.52 M, 2.76 M, and 1.38 M. It is to note, however, that the experiments did not cover residue and practical application aspects, and further extensive studies are needed in this area to clarify the relevance of the practical sub-classification.

## Introduction

The two-spotted spider mite, *Tetranychus urticae* (C.L. Koch, 1836) (*Arachnida: Tetranychidae*), is a cosmopolitan polyphagous mite species. It can cause significant qualitative damage and yield losses in horticultural and arable crops, as well as in greenhouses and orchards (Stumpf et al., 2001, Van Leeuwen et al., 2007). Both larva, nymphs and adult developmental stages feed mainly on the lower leaf surface of plants. Their damage can be reflected in reduced photosynthesis and the introduction of phytotoxic substances into the plant through the irreversible destruction of the inner plant structures. (Johnson and Lyon 1976). In addition, web formation, mite excrement, and defoliation can affect the plant’s external appearance, thereby reducing its commercial value (Johnson and Lyon 1976). Spider mites might be vectors of various plant viruses, e.g., potato Y virus (Schulz 1963). However, subsequent studies have not confirmed it (Orlob 1968).

Its hidden lifestyle and high reproductive and dispersal capacity impose severe constraints on its defence. The residual pesticides on the market used to control mites are limited (Leeuwen et al., 2009). Moreover, the species can develop a resistance to miticides relatively quickly (Vrie et al., 1972, Leeuwen et al., 2009). Chemical control agents also pose a significant environmental burden and can adversely affect humans and even some beneficial organisms. There is an ongoing need for new miticide compounds to provide for the successful realisation of sustainable and environmentally friendly management against it.

Since synthetic miticides are not always sufficiently effective to keep mites below the damage threshold, plant extract-based pesticides are emerging as a promising option (Heuskin et al., 2011). Using mixtures of biosynthetic compounds (mineral oil, entomopathogen fungi, essential oil, kaolin particle) based on plant extracts and practical application of some natural enemies as biopesticides can restrain the development of resistance (Isman 2000). These products are biodegradable with less environmental impact and non-desired side effects (Hay and Waterman 1993, Chiasson et al., 2001, Isman 2001, Basta and Spooner-Hart 2002, Isman 2004). Nevertheless, their application may be limited by the problem of manufacturing them in the required quantities and in a sustainable way (Isman 2000). One of these natural compounds is lithium chloride, which has been proven effective against *Varroa destructor* (Anderson & Trueman, 2000), the most important parasite of *Apis mellifera* (Linnaeus, 1758). The efficacy of lithium chloride has not been examined in other important pests yet. Like the spider mite, *V. destructor* is problematic to control because of its hidden lifestyle and the development of resistance. Studies on honey bees indicate that lithium chloride has considerable potential for use in beekeeping (Kolics et al., 2021). The aims of our study were to obtain information about the efficacy of the lithium chloride against the *T. urticae*, and to evaluate the influence of the dose rate of this potential active substance. By means of these initial results, we are going to enrich the plant protection knowledge of this serious agricultural pest, giving an opportunity for the realisation of a successful control method against it.

## Materials and methods

### Experimental design

*T. urticae* specimens of mixed age, sex and gender were collected from non-spraying soybean fields in three places at Kaposvár, Hungary (N: 46.39 E: 17.85) in June 2021. Three subsamples of *T. urticae* were collected from five plant leaves from each location. Sampling areas were separated by at least 1,000 m, and 300-500 spider mites were collected in each subsample. *T. urticae* mixed and transferred to host soybean plants; these plants were not treated with any zoocide previously. The mites were kept on the plant until they were used in the experiments under laboratory conditions (at 25±1°C, 60±5% relative humidity (RH), and a photoperiod of 16:8 (L:D) h each day. Altogether 60 individuals were investigated in each treatment (20 individuals in 3 replications).

### Acaricidal bioassay

The contact micro-immersion bioassay was performed according to Dennehy et al. (Dennehy et al., 1993) with slight modifications as each batch of 20 mites was transferred with a pipette tip to immerse them into a 1.5 ml Eppendorf tube containing 1 ml aqueous lithium chloride solution of the following concentrations: 5.52 M, 2.76 M, and 1.38 M (corresponding to those applied in an earlier experiment carried out against the mite parasite of the honey bee (Kolics et al., 2020)).

Subsequently, they were placed on a filter disc (Sartorius, d=150mm, Grade: 1292) in a petri dish and surrounded with wet cotton wool. To preselect individuals, mites that did not show vigorous movements were discarded, leaving 20 vital individuals on each dish, and their activities were studied using a binocular microscope. Each treatment was performed in three iterations, the control treatment was also carried out with an aqueous solution.

The first recorded sign of lithium poisoning was regarded at the onset of uncontrolled tremorous locomotion performed by the first mite of the block of 20 individuals. Subsequently, the occurrence of the first dead mite in the block, the occurrence of 50% dead mites, the occurrence of 90% dead mites, and the occurrence of all dead mites in the block was recorded. Each bioassay was replicated three times.

### Statistical analysis

The elapsed times from the beginning of the treatments were analysed separately for each treatment for the recorded events detailed above. Abbott’s formula (Abbott 1925) was applied to compute mortality rates. Data were transformed by the ln transformation (natural logarithm). The Kolmogorov-Smirnov and the Shapiro-Wilk tests carried out normality testing for the above events. All events were found to be normally distributed (p>0.05). Statistical differences were analysed with one-way ANOVA for each event separately. The Levene test justified the homogeneity of variances (p>0.05). Pairwise differences were analysed with Tukey’s post hoc test. The mortality rates computed in the statistical tests were calculated by SPSS 22.0 software.

## Results

Abbott’s corrected mortalities of *T. urticae* experimental population triggered by lithium chloride are shown in Fig.1. and Table 1. Depending on the exposure times, the mortality character of some doses was logarithmic type in any case. Mortalities were unequivocally increased by the exposure times. Besides, their tendencies were approached to the total eradication degree by a steeper curve and relatively faster. The symptoms induced by the treatment were similar to those observed in *V. destructor*, as uncontrolled tremorous insect movement was followed by death, except that the immobility phase was not observed in the case of the two-spotted spider mite. Recovery from paralysis was observed on some mites in the 0.69M concentration. Therefore, observation of LT90 was discontinued.

**Figure 1.**
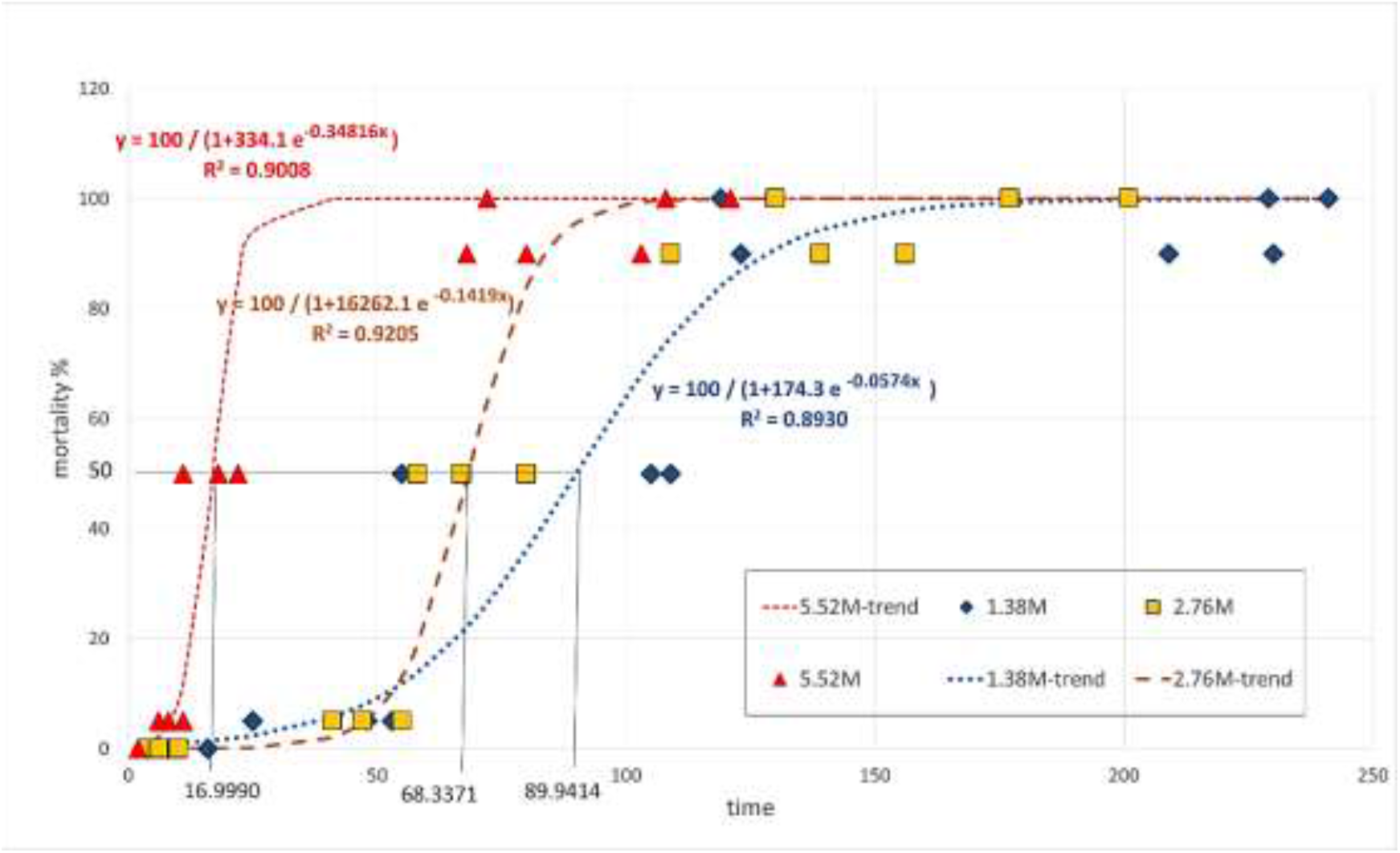
Abbott corrected mortality rates for the three treatments

**Table 1.** Mean ± SE (standard error) values for the occurrence of events by treatment

| Treat-<br>ment | 1<sup>st</sup> Symptom | 1<sup>st</sup> Dead | LD<sub>50</sub> | LD<sub>90</sub> | LD<sub>100</sub> |
| --- | --- | --- | --- | --- | --- |
| 1.38M | 9.33 ± 3.528 | 42.00 ± 8.622 | 89.6 7± 17.372 | 187.33 ± 32.733 | 196.33 ± 38.822 |
| 2.76M | 6.67 ± 1.764 | 47.67 ± 4.055 | 68.33 ± 6.386 | 134.67 ± 13.740 | 169.33 ± 20.851 |
| 5.52M | 1.67±0.333 | 8.33 ± 1.453 | 17.00 ± 3.215 | 83.67 ± 10.269 | 100.33 ± 14.655 |
| The trend of mortality rates (y) by exposure times (x) |  |  |  |  |  |
| 1.38M | $y = 100 / (1 + 16262.1 \times e^{-0.1419x})$; $R^2 = 0.9205$ | | | | |
| 2.76M | $y = 100 / (1 + 174.3 \times e^{-0.0574x})$; $R^2 = 0.8930$ | | | | |
| 5.52M | $y = 100 / (1 + 334.1 \times e^{-0.34816x})$; $R^2 = 0.9008$ | | | | |

As is shown in Figure 1, a classic logistic growth curve (also known as the Pearl-Reed logistic curve) was fitted to the Abbott corrected mortality rate data (Abbott, 1925) for each treatment, of the form y = K / (1 + b × e-cx) with K=100, and y representing the mortality rate and x the exposure time. The fitted equations are shown in Figure 1 and Table 1, with the respective R^2^ values (greater than 0.89).

Figure 2 gives the needed exposure times for reaching the LD_50_ stage by dosage level. The exposure times show a definite decreasing trend with higher dosages and the fitted quadratic trend. As there are only three dosage levels, it is not very reliable to extrapolate. Still, by simple derivation of the trend equation, the rate of decrease is 150.2 x (dosage level) along the curve.

**Figure 2.**
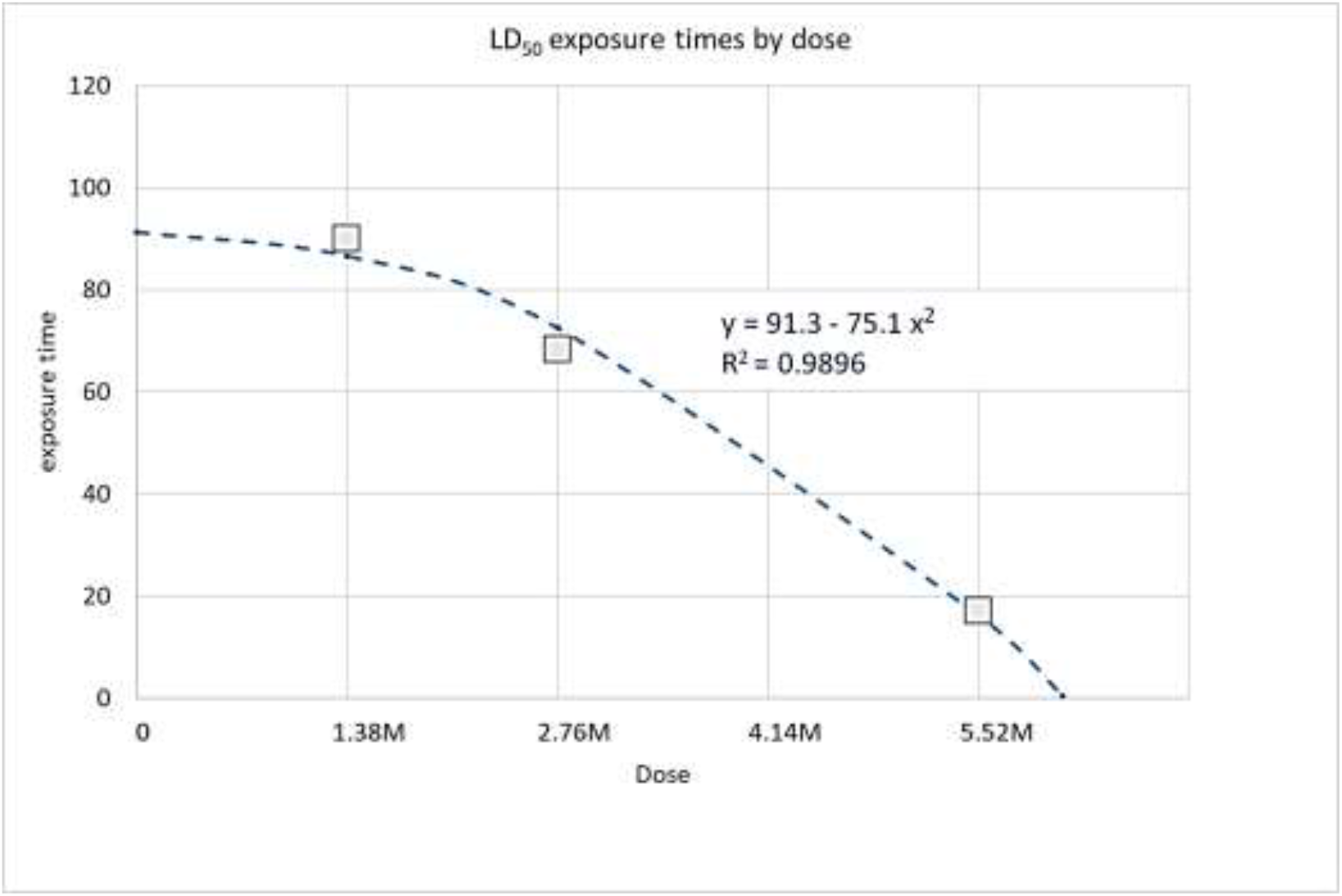
Estimated exposure times to LD50 by dosage

Table 1 gives the observed mean exposure times needed to reach the given mortality stages for each treatment. As shown in Table 1 and Figure 3, the largest dose required only half of the exposure time of the smallest dose for the LD100 stage, while the difference is even greater for LD90, especially LD50. Similar tendencies hold for the other mortality stages, too. The standard errors (SE) in Table 1 and the confidence intervals in Figure 3 also indicate that the largest dosage produced the most stable exposure times having the smallest SE values for any stage. This is supported by the parameters of the fitted logistic equations for the three treatments, presented in Table 1, too.

**Figure 3.**
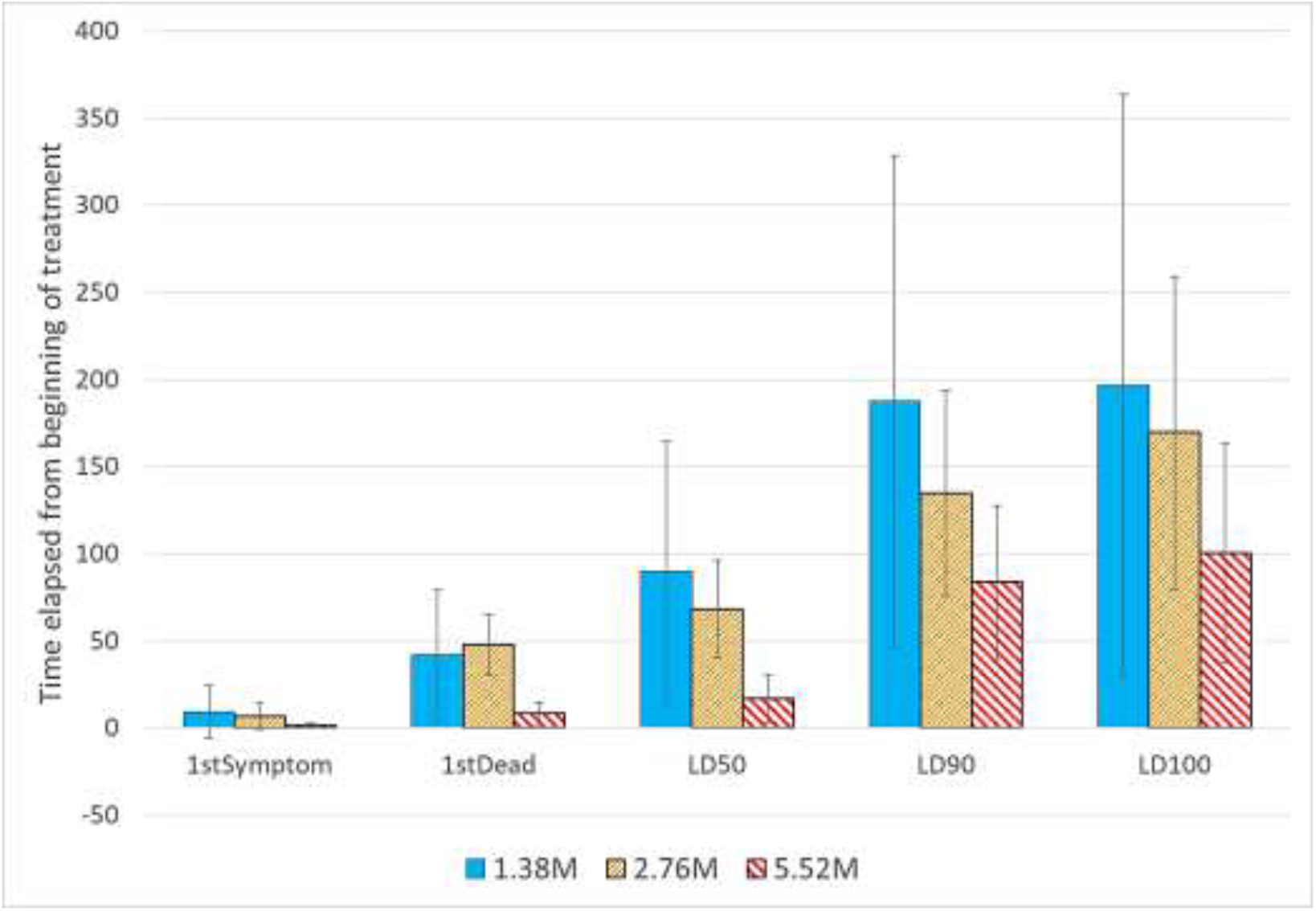
Average exposure times by treatments, with 95% confidence intervals for the mortality stages

One-way ANOVA results are presented in Table 2 to test the significant differences between the mean ln-transformed exposure times of the three treatments with three replications for each treatment. As the last column of Table 2 indicates, treatments significantly differ at p=0.05 in the occurrence times of each mortality stage, except the last one (LD100), where only at p=0.1 can significant differences be accepted.

**Table 2.**
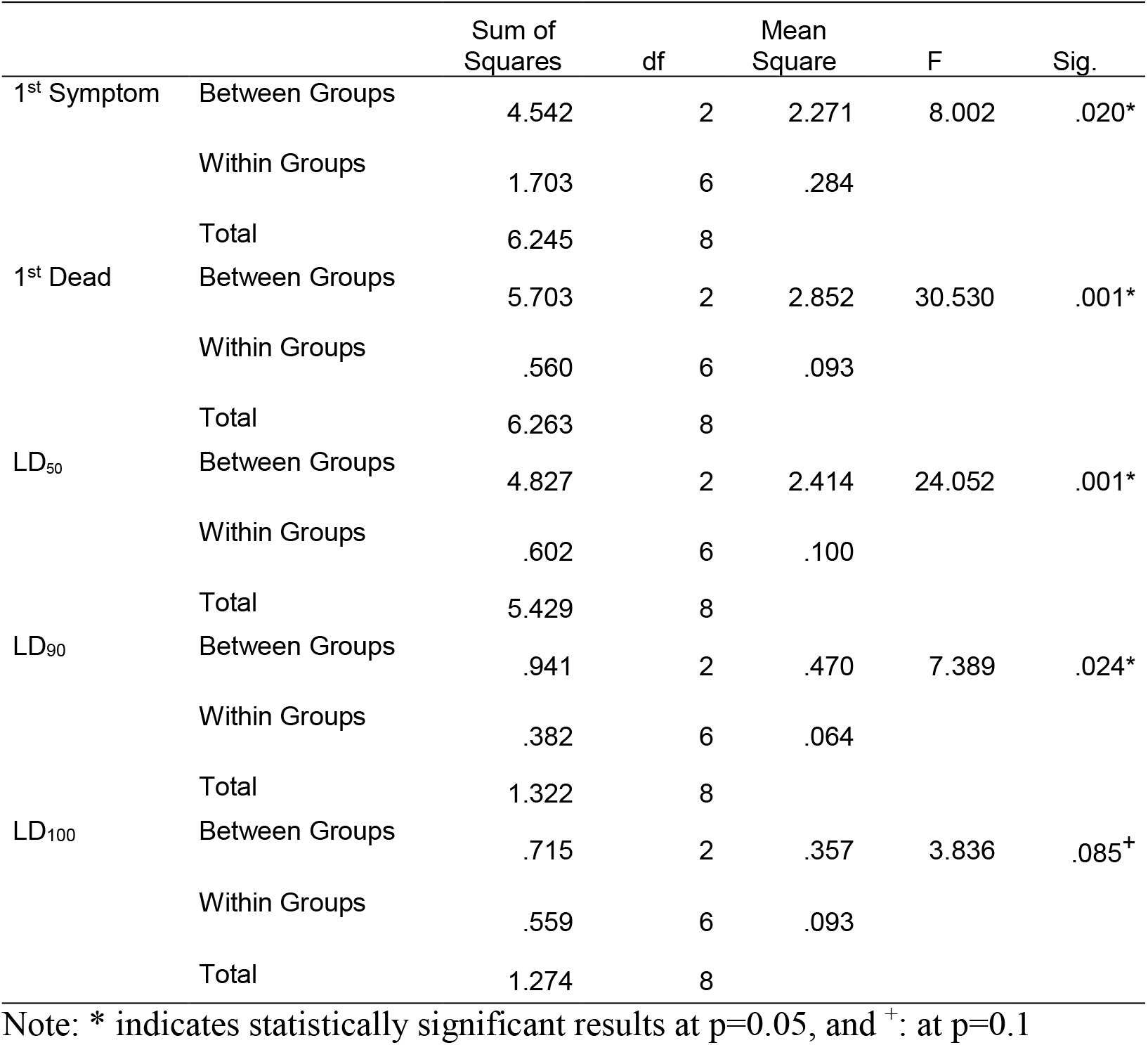
One-way ANOVA for the natural logarithms of the occurrence times by treatment

A post hoc test (Tukey HSD) was applied to compare the treatments pairwise with regard to exposure times to the mortality stages. As Table 3 shows, the largest dosage significantly differs from the other two treatments up to the occurrence of 50% dead mites (LD50), and it significantly differs from the smallest concentration for the occurrence of 90% dead mites (LD90). However, treatments do not differ significantly in the occurrence of 100% dead mites.

**Table 3.** Tukey (HSD) test results for separating subsets (alpha = 0.05) - mean values of subsets for the logarithms of occurrence times

|  | 1<sup>st</sup> Symptom |  | 1<sup>st</sup> Dead |  | LD<sub>50</sub> |  | LD<sub>90</sub> |  | LD<sub>90</sub> |
| --- | --- | --- | --- | --- | --- | --- | --- | --- | --- |
| treatment | subset |  | subset |  | subset |  | subset |  | subset |
|  | 1 | 2 | 1 | 2 | 1 | 2 | 1 | 2 | 1 |
| 5.52M | 0.4621 |  | 2.0897 |  | 2.7931 |  | 4.4121 |  | 4.5849 |
| 2.76M |  | 1.8269 |  | 3.6868 |  | 4.2157 | 4.8919 | 4.8919 | 5.1157 |
| 1.38M |  | 2.0794 |  | 3.8570 |  | 4.4509 |  | 5.1975 | 5.2325 |
| Sig. (p) | 1.000 | 0.835 | 1.000 | 0.782 | 1.000 | 0.655 | 0.127 | 0.362 | 0.090 |

## Discussion

Our results highlight the potential of lithium salts as a control alternative for other mite species, such as the two-spotted spider mite. These results are also significant because the control of *T. urticae* is complicated in pest management due to the species’ aggressiveness, polyphagous and cosmopolitan nature. In recent years, interest in nature-identical pesticides has increased significantly due to environmental concerns and the resistance of the two-spotted mites to conventional pesticides.

Alkali metal lithium is widely distributed in the environment, the 27th most abundant element of the earth’s crust (Habashi 1997). Lithium-ion is not biodegradable; however, it is not expected to bioaccumulate. Pollution problems are mostly related to cases where lithium is disposed of with heavy metals (Aral and Vecchio-Sadus 2008). Neither lithium intake from food and water nor occupational exposure presents a toxicological hazard. It does not appear to pose a considerable threat to flora and fauna on land or water (Aral and Vecchio-Sadus 2008). It can be a potential natural micronutrient component of foodstuffs, with its quantity depending mainly on the soil microelement lithium content in amounts of the order of a milligram. (Bogdanov et al., 2008, Tutun et al., 2019).

Although in large quantities, it can be toxic to plants, animals and humans, with significant differences in susceptibility depending on the organism (Shahzad et al., 2016). On the other hand, lithium is reported to have rather beneficial health effects, as studies indicate lower suicide rates in populations that consume water with higher ionic Li content (Schrauzer and Shrestha 1990, Ohgami et al., 2009, Kapusta et al., 2011, Giotakos et al., 2013, Sugawara et al., 2013, Liaugaudaite et al., 2017). Moreover, in recent years, modern psychiatry has proposed the production of foods fortified with lithium, similar to the model of iodisation of table salt (Marshall 2015, Terao 2015, TM 2015), in agreement with former findings (Schrauzer 2002). Its mode of action or role is disputed; intracellular accumulation of Li results in the replacement of Sodium, which in turn reduces intracellular Ca^2+^ concentration, inhibits release, and facilitates the uptake of major transmitters: noradrenaline, serotonin, and dopamine (Won and Kim 2017). The veterinary potential of lithium appeared as the most recent feature of the trace element. The potential use of lithium salts in beekeeping is approaching technology-level application (Stanimirovic et al., 2019, Kolics et al., 2021). Nonsynonymous mutations in the pesticide target site of synthetic acaricides are often reported (Feyereisen et al., 2015), implying an inevitable risk of long-term resistance.

Although there are similarities between the use of alkaline earth metals and the use of copper compounds, which are also commonly used, several publications have discussed how their longterm use alters the mineral composition of the soil (Brun et al., 2001). On the other hand, lithium is not heavy metal, unlike copper. Moreover, it is a mobile element (Aral and Vecchio-Sadus 2008). Supposing its vegetative parts remain in the area, the potential use of lithium may increase its quantity in soil compared to its original presence. Although it is encouraging that there seem to be some species of mites, such as the *Thyrophagus putescentiae* (Schrank, 1781) species we studied, on which lithium does not show efficacy, it is also essential to study the effect of lithium on parasitic mite species (e.g. *Phytoseiulus sp*., *Ambliseius sp*.) used in certain crops to reveal the risks of its potential use.

Our studies were limited to in vitro experiments and did not include the lower concentrations used for *Varroa* mites. Half of the lowest concentrations we tested already had individuals recovering from poisoning at 1/8 concentration. It is to note that concentrations below 0.69 M may also exert satisfactory miticidal effectiveness that may be enhanced using surfactants. The utilisation of lithium chloride can be an effective solution against *T. urticae* in several crops, and its rational use can reduce pest populations in sunflower, soybean, maise, or vineyard. In this way, studying the possibilities of a potential lithium-based chemical control technique is justified for further investigation in the quest for the satisfying mapping of all environmental consequences.

## Acknowledgements

This publication and the APC were supported by the Hungarian Government and the European Union, with the co-funding of the European Regional Development Fund in the frame of Széchenyi 2020 Programme GINOP-2.3.2-15-2016-00054 project.

